# Mutualism between microbial populations in structured environments: the role of geometry in diffusive exchanges

**DOI:** 10.1101/172924

**Authors:** François J. Peaudecerf, Freddy Bunbury, Vaibhav Bhardwaj, Martin A. Bees, Alison G. Smith, Raymond E. Goldstein, Ottavio A. Croze

## Abstract

The exchange of diffusive metabolites is known to control the spatial patterns formed by microbial populations, as revealed by recent studies in the laboratory. However, the matrices used, such as agarose pads, lack the structured geometry of many natural microbial habitats, including in the soil or on the surfaces of plants or animals. Here we address the important question of how such geometry may control diffusive exchanges and microbial interaction. We model mathematically mutualistic interactions within a minimal unit of structure: two growing reservoirs linked by a diffusive channel through which metabolites are exchanged. The model is applied to study a synthetic mutualism, experimentally parameterised on a model algal-bacterial co-culture. Analytical and numerical solutions of the model predict conditions for the successful establishment of remote mutualisms, and how this depends, often counterintutively, on diffusion geometry. We connect our findings to understanding complex behaviour in synthetic and naturally occurring microbial communities.

## 1 Introduction

Microorganisms display a broad spectrum of interactions that determine the behaviour of microbial communities (Abreu and Taga 2016). Predicting this behaviour is a fundamental challenge in current microbial ecology (Widder et al. 2016). A wealth of experimental data on microbial community structure and dynamics is now available from ‘omics’ approaches (Cooper and Smith 2015, Widder et al. 2016). These, however, need to be complemented by lab-based studies of synthetic consortia and mathematical models to reach a mechanistic understanding of microbial dynamics (Widder et al. 2016, Abreu and Taga 2016). The study of mutualistic interactions between microbial populations is an active area of current resesarch in microbial ecology. Recent experimental studies have investigated synthetic mutualisms between microbes across the kindoms of life. These include strains of enteric bacteria (Harcombe et al. 2014, McCully et al. 2017, LaSarre et al. 2017) and yeast (Allen et al. 2013) engineered to be mutualistic, and synthetic consortia combining wild type microbial species, such as bacterial tricultures (Kim et al. 2008), mixed cultures of algae and fungi (Hom and Murray 2014), and algae and bacteria (Croft et al. 2005, Kazamia et al. 2012, Wang et al. 2014, Segev et al. 2016).

Mutualistic interactions are conventionally modelled using Lotka-Volterra type models, with positive interaction coefficients (Murray 1989). Linear mutualistic Lotka-Volterra models are known to display unrealistic unbounded growth (Murray 1989), but logistic versions have been used to study demographically open mutualistic populations (Thompson et al. 2006), transitions between interspecies interactions (Holland and DeAngelis 2009, 2010), and the steady state dynamics of algal-bacterial co-cultures (Grant et al. 2014). Since the pioneering work of May (1973), such models have also been fruitfully employed to describe mutualistic interactions in network models of communities (Okuyama and Holland 2008). In such models the interaction coefficients coupling species together define an interaction or community matrix (for mutualistic interactions the coefficients are positive and symmetric). Significant shortcomings of Lotka-Volterra models have recently been pointed out. For example, when species interact by exchanging metabolites, a metabolite-explicit model does not in general map onto a Lotka-Volterra implicit model (Momeni et al. 2017). Only in special instances does the microbial Lotka-Volterra form provide a good description of the microbial dynamics, e.g. when a fast equilibration approximation holds (Hoek et al. 2016). Resource-explicit models of bacterial mutualisms compare well with experiments in which mutualists are *well-mixed* (Lee et al. 1976, Wang et al. 2007, LaSarre et al. 2017, McCully et al. 2017). Explicitly modelling resources is critical when studying spatially structured mutualistic systems (not well-mixed) whose interactions are contolled by metabolite dynamics and their spatial transport.

Recent studies have considered spatial aspects of mutualistic and cooperative microbial interactions. Simulations using flux balance analysis (FBA) successfully predict the spatial growth on agar of colonies of synthetically mutualistic enteric bacteria (Harcombe et al. 2014). The FBA approach requires explicit knowledge of every known metabolic biochemical pathway in each mutualistic species, restricting its applicability to mutualisms between metabolically well-characterised organisms. Spatial effects on cheating (Momeni et al. 2013) and genetic drift (Müller et al. 2014) observed in yeast colonies growing on agarose pads have also been modelled explicitly. In these models, coupled cells and nutrients diffusing in two dimensions are simulated to predict how nutrient-mediated interactions control spatial heterogeneity and survival of the populations. In general, interactions have been shown to control the spatial structure of laboratory biofilm communities (Nadell et al. 2016). However, the homogenous environment of nutrient agarose or laboratory biofilm substrates do not possess the intrinsic geometric or topological structure of natural microbial environments, such as the porous matrix of soil or microfluidic analogues (Coyte et al. 2017). Mutualistic microbial dynamics have not thus far been studied in such structured environments, to the best of our knowledge.

Here, we study a model of mutualistic microbial species in a simple geometry representing a minimal unit for a structured environment: populations growing in spatially separated reservoirs, metabolically linked by a channel. As soil can be approximated at the microbial scale as a physical network of growth chambers linked by channels (Pérez-Reche et al. 2012), this model provides the basis for the description of natural complex porous networks or artificial regular networks (see Figure 1) and the microbial interactions therein. The model is generally applicable to auxotrophs cross-feeding remotely. We apply it to make predictions for the dynamics of mutualistic populations of algae and bacteria exchanging vitamin B_12_ and a carbon source (Kazamia et al. 2012). Our predictions provide new insights into the behaviour of microbial communities residing in structured geometries, both within synthetic consortia in the laboratory and environmental microbial communities.

**Figure 1:**
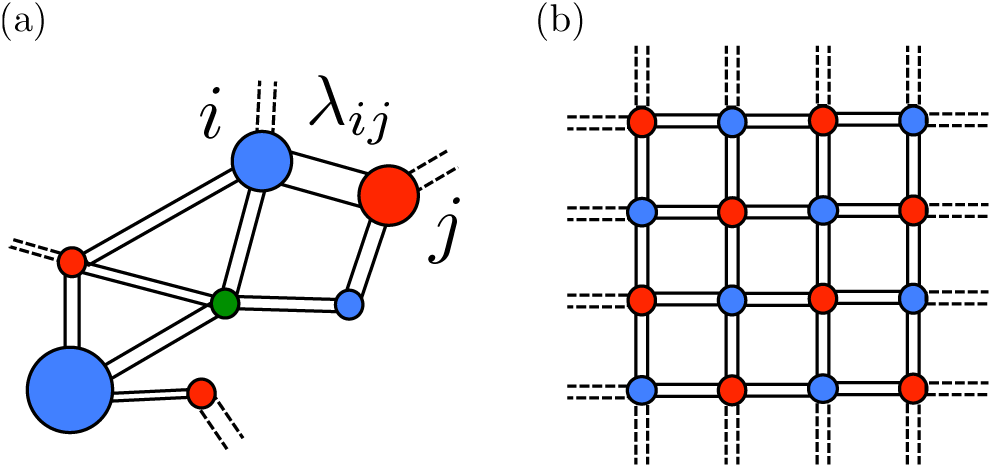
Schematic of a diffusively coupled microbial network representing: (a) A structurally and microbially heterogeneous network as a realistic representation of soil (Pérez-Reche et al. 2012); (b) A crystalline network that can be engineered in the laboratory. The nodes of this physically structured network represent reservoirs of different volumes filled with different growing microbial species diffusively exchanging metabolites via porous channels, as described in the model formulated in this work. Diffusive exchanges are parameterised by sets of geometric parameters, as such as the lengths, *λ*_*ij*_, of the channels connecting nodes.

## 2 Model

The model describes two populations of mutualistic microbial species, A and B, interacting at a distance. The mutualistic interactions are predicated on auxotrophy: A requires metabolite V (for “vitamin”), excreted by B; conversely B requires metabolite C (for “carbon”), excreted by A. In formulating the problem we shall first use variables with an overbar to denote dimensional quantities (concentrations, time, space), reserving symbols without typographical modification for appropriately rescaled variables. Populations of A and B, with densities 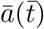 and 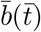 respectively, reside in two well-mixed reservoirs, of equal volume Γ. These are spatially separated, but connected by a cylindrical channel (length *L*, cross-sectional area Σ), as in Figure 2. The channel is impervious to cells, but porous to metabolite exchange by diffusion. Population A produces metabolite C with local concentration 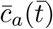, which diffuses out of the reservoir and into the channel at 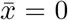 (with 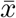 denoting the position along the channel axis), where it develops a spatial profile 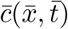 and eventually reaches the other reservoir at 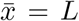, where its concentration is 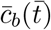. Symmetrically, metabolite V produced by B with concentration 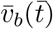, diffuses out at 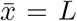 giving 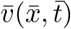, feeding the other reservoir at 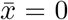, generating a concentration 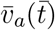.

**Figure 2:**
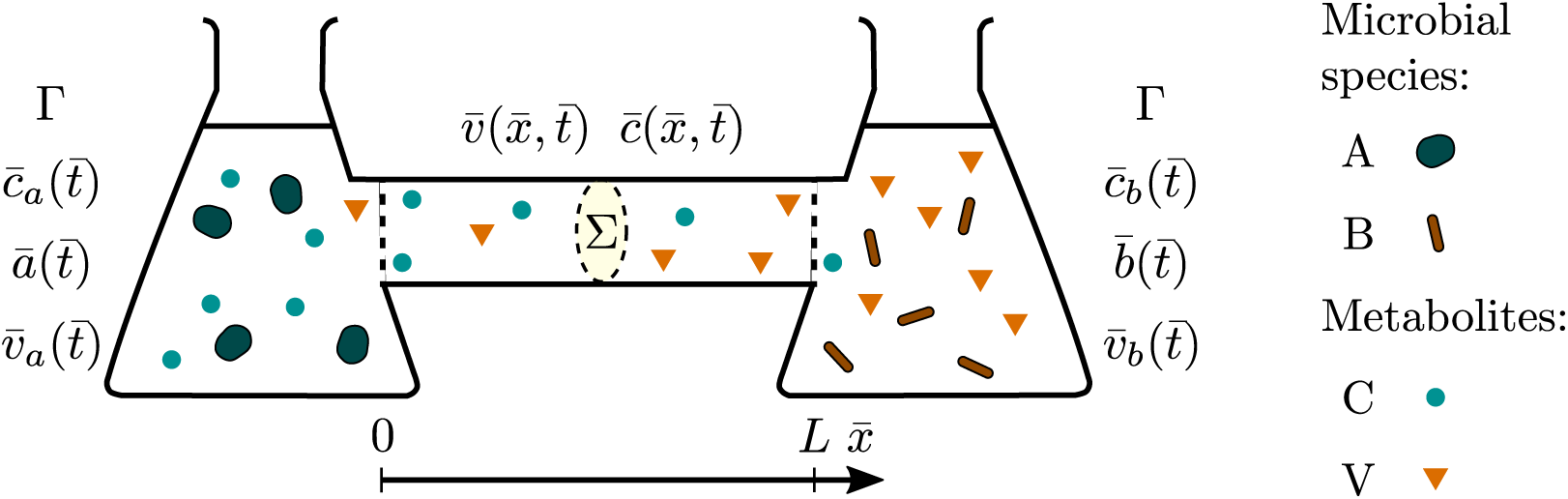
Diffusive cross-feeding at a distance. Auxotrophic microbial populations A and B (concentrations 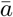 and 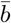) reside in well-mixed reservoirs of equal volume Γ separated by a channel of length *L* and cross-section Σ. Microbe A produces a carbon source C, of homogeneous concentration 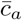, in its reservoir. This diffuses through the channel, forming a profile 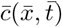, a function of position along the channel 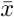 and time 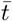. On reaching the reservoir where microbe B resides the concentration is homogenised to 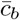. Symmetrically, the vitamin V produced by microbe B in its reservoir at concentration 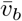, diffuses to reservoir A creating a profile 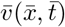, homogenised to 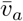 in the reservoir. Here, this general model is applied to an algal-bacterial partnership.

We first consider dynamics within the channel connecting the reservoirs, within which metabolites obey one-dimensional diffusion equations,

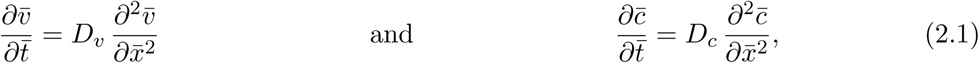

with *D*_*s*_ the diffusion coefficients for metabolite S = C or V. The boundary conditions to (2.1) obtained from continuity at the channel-reservoir interface are: 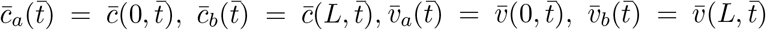. Clearly, one characteristic time scale of the problem is set by diffusive equilibration along the length of the channel,

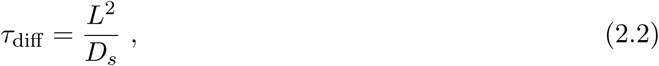

where we anticipate that the diffusion constants of both metabolite species are similar. From Fick’s law, the flux *J*_*s*_ (molecules area^−1^ time^−1^) of metabolite species S (C or V) entering, say, the left reservoir from the channel is

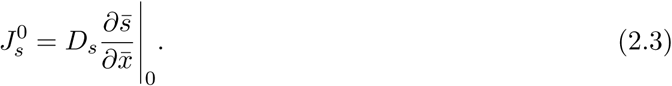

The rate such molecules enter the reservoir is 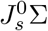, and with instantaneous homogenisation there, the rate of change of the reservoir concentration 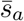 is 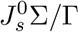. The characteristic length

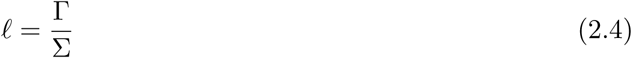

will play an important role in the model. If 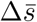 is a typical difference in concentration of S between the two reservoirs, then the typical gradient within the channel is 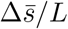, giving rise, by the arguments above, to an associated rate of change of reservoir concentration scaling as 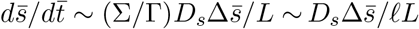, from which we can identify a characteristic equilibration time

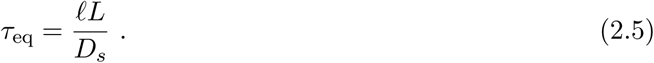

We define the ratio of equilibration and diffusive time scales to be

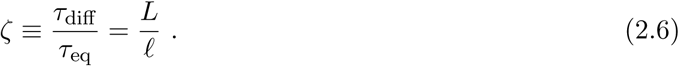

The regime *ζ* ≪ 1 is that of fast establishment of the linear concentration profile in the tube relative to changes of concentrations in the reservoirs, while for *ζ* ≥ 1 the transients within the channel are on comparable time scales to that for changes in the reservoirs. Semi-analytical solutions to the problem of chemical diffusion between two connected reservoirs further demonstrate the existence of these two regimes and the role of the previously identified timescales (see Supporting Information).

We now turn to the population dynamics within the reservoirs, in which we explicitly assume that algae reside in reservoir A and bacteria in B, and that vitamin B_12_ and carbon are exchanged. The dynamics obey the ordinary differential equations

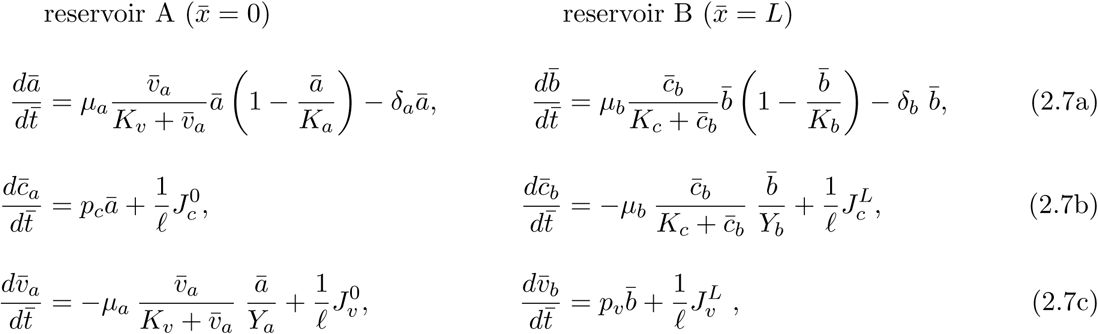

where 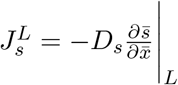 is the flux of metabolite S = C or V entering the right reservoir. In equations (2.7a) we model cell growth as logistic, with maximum growth rate *µ*_*i*_ and carrying capacity *K*_*i*_ for species *i* = A or B. Growth rates are limited by the abundance of the required metabolites. This is modelled using Monod factors (Monod 1949), e.g., for C, 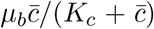, where *K*_*c*_ is the half-saturation constant (and symmetrically for V). Linear death terms, with mortality rates *δ*_*i*_ for *i* = A or B, ensure exponential negative growth in the absence of the limiting metabolites. Equations (2.7b) describe the dynamics of metabolite C. This is produced by species A in proportion to its concentration with a rate *p*_*c*_, and diffuses out at 0. In the other reservoir, C is taken up by B. The uptake is assumed proportional to the cell growth rate, the proportionality constant is 1*/Y*_*b*_, where *Y*_*b*_ is the yield coefficient (how much metabolite C results in a given concentration of species B). Equations (2.7c) describe the V dynamics, which are completely symmetric to the C dynamics. Although inspired by bacterial-algal symbiosis, it is clear that the structure of these dynamics is quite broadly applicable to mutualistic systems in general.

### Identifying the key model parameters

In order to access the general dynamics of remotely cross-feeding monocultures, we nondimensionalise equations (2.7). Because our focus is on the impact of geometry on the biological processes, we choose a scheme accordingly. First, normalize the bacterial and algal concentrations by their respective carrying capacities, the organic carbon and vitamin concentrations by their respective half-saturation concentrations, rescale time by the bacterial growth rate, and rescale space by the length scale 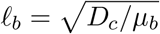 of organic carbon diffusion on the time scale of bacterial growth, defining

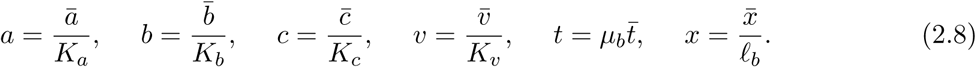

The ratios of algal and bacterial growth rates and of their diffusion constants,

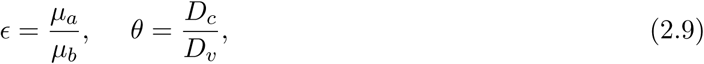

are two additional parameters. With now three characteristic lengths in the problem (*L, l, l*_*b*_) one can form two independent dimensionless ratios. These can be taken to be

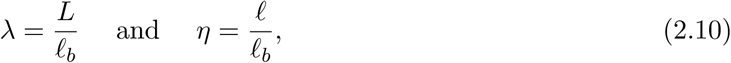

so that the parameter *ζ*, defined previously in Eq. 2.6, is *ζ* = *λ/η*.

There are three pairs of parameters remaining which capture the relative strength of cellular death, uptake and production in bacteria and algae respectively. They are: the ratios of death rate to maximum growth rate of bacteria and algae, which define mortality parameters

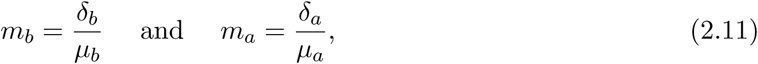

which must be less than 1 for any population increase to occur; and finally, for both carbon and vitamin, the ratios of the typical uptake rate to the typical rate of change define the uptake parameters

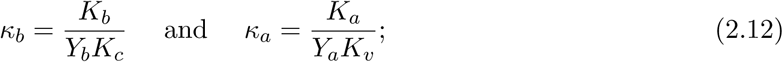

for both carbon and vitamin, the ratios of the typical production rate to the typical rate of change define the production strengths

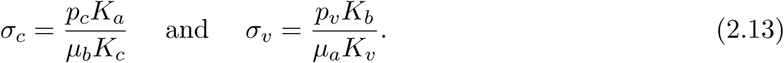

With these rescalings, the dimensionless evolution equations are

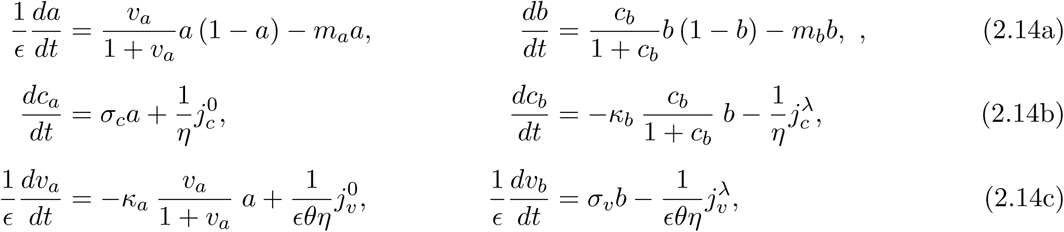

where now the dimensionless fluxes are 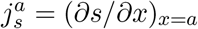. These equations are to be solved together with the diffusion equations

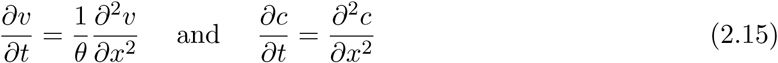

for *c* and *v* on the interval *x* ∈ [0, λ], ensuring continuity of fluxes and concentrations at the ends of the tube. Equations (2.14) were solved numerically to explore the role of diffusive geometry on mutualistic coexistence. We used the nondimensional parameters shown in Table 1, corresponding to the mutualistic association between *Lobomonas rostrata*, a B_12_-requiring green alga, and *Mesorhizobium loti*, a B_12_-producing soil bacterium (Kazamia et al. 2012). Parameter values were obtained by fitting growth and vitamin B_12_ assay data from co-culture experiments with this synthetic mutualism (see Supporting Information and Figure A.2).

**Table 1:** Non-dimensional model parameters for the mutualistic association of *M. loti* and *L. rostrata*.

| Non-dimensional parameter | Symbol | Value |
| --- | --- | --- |
| <u>Biological parameters</u> |  |  |
| Uptake parameter for algae | $\kappa_a$ | 1.3 |
| Uptake parameter for bacteria | $\kappa_b$ | 2.2 |
| Algal mortality/growth ratio | $m_a$ | 0.024 |
| Bacterial mortality/growth ratio | $m_b$ | 0.014 |
| Carbon production strength | $s_c$ | 0.018 |
| Vitamin production strength | $s_v$ | 3.2 |
| Algal to bacterial growth rate ratio | $\epsilon$ | 0.72 |
| <u>Physical parameters</u> |  |  |
| Ratio of metabolite diffusivities <sup>a</sup> | $\theta$ | 2.5 |
| Channel length | $\lambda$ | 1–30 |
| Equilibration length | $\eta$ | 3–100 |

<sup>a</sup> obtained considering carbon with diffusivity $D_c = 5 \times 10^{-6} \text{ cm}^2 \text{ s}^{-1}$ as metabolite C and B<sub>12</sub> vitamin with diffusivity $D_v = 2 \times 10^{-6} \text{ cm}^2 \text{ s}^{-1}$ as metabolite V

Before discussing the results from numerical solutions of the dynamical system of our model, we note that it exhibits a set of fixed points for which the cell concentrations vanish (*a* = *b* = 0) at any combination of residual metabolite concentrations. More interestingly, a single positive fixed point exists given by equations (A.7) below under the conditions of equations (A.8) (see Supporting Information). This positive fixed point corresponds to an established co-culture at a distance.

### Feeding on a distant passive source

Before considering the fully coupled system dynamics, we consider the case of a single auxotrophic species B, concentration *b*, residing in a reservoir initially free of a growth-limiting metabolite coupled by the channel (also initially nutrient-free) to a strong source of the metabolite with initial concentration 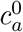. This source consists of a reservoir filled with limiting metabolite. The long time steady-state for the model is always extinction of B once it has exhausted the remote resource. However, separation of the microbial population from the source modifies the transient population dynamics. Recalling the nondimensional channel length 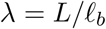 and equilibration length 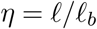, we can define the nondimensional timescales *t*_diff_ = λ^2^ and *t*_eq_ = *λη* as the ratios between the typical times of diffusion and of equilibration between reservoirs, and the biological growth timescale *τ*_b_ = 1*/µ*_*b*_. These ratios gauge the relative rates of diffusion/equilibration and growth. We require *t*_eq_ and *t*_diff_ *~* 1 for diffusion to transport metabolites to species B, stimulating its growth.

We have solved the remotely-fed single microbe limit of the model (see Supporting Information) to predict the dynamics of the rhizobial bacterium *Mesorhizobium loti* fed from a remote glycerol carbon source. Figure 3 shows the transient growth dynamics in the regime for which both geometric parameters *λ* and *η* impact the dynamics. We first consider the effect of diffusive reservoir equilibration, quantified by *η* for a fixed channel length *λ*. For large *η*, *t*_eq_ is large: diffusive equilibration in the reservoir is much slower than growth. Thus, the instantaneous flux from the carbon source reservoir to the bacterial reservoir is below what the bacteria need to grow to carrying capacity. As a result, increasing *η* decreases the value of the peak bacterial concentration (preceding the inevitable decay), as well as delaying the onset of growth (Figure 3a). Next we fix *η* and vary *λ*. Since the diffusive timescale scales like *λ*^2^, increasing *λ* progressively delays the onset of bacterial growth (Figure 3b inset). Large *λ* values also correspond to weaker carbon source gradients across the tube, and thus a ‘slow-release’ nutrient flux. Consequently, a less concentrated population can be sustained for longer by the remote source (Figure 3b). The passive source case we have just considered demonstrates the critical role played by both geometric parameters *λ* and *η* in setting the timescale of transients, but also the peak microbial numbers achievable on a finite resource.

**Figure 3:**
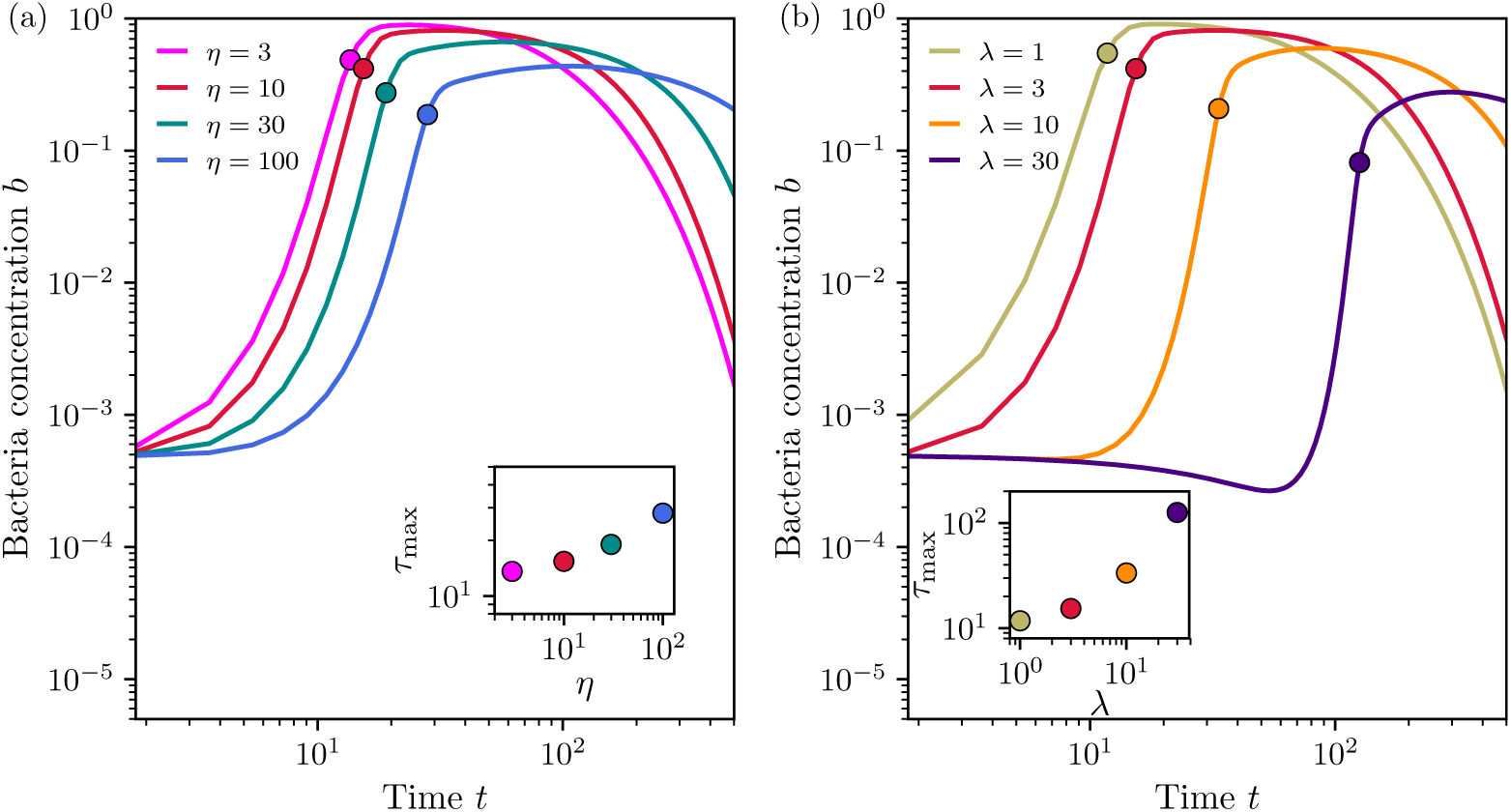
Transient dynamics of a bacterial population fed through a channel that allows metabolite diffusion from a remote carbon source. The diffusive exchange geometry controls the dynamics through the nondimensional channel length *λ* and reservoir equilibration length *η*. Model solutions predict that: (a) for fixed *λ* = 3, increasing *η* delays the time of peak bacterial growth and curtails growth due to a limited carbon-source flux; (b) for fixed *η* = 10, increasing *λ* significantly delays peak growth, with an impact on the maximum bacterial concentration attained. The delay as measured by *τ*_max_, the time of maximal growth rate, is proportional to *λ*^2^ (inset). For all simulations, initial nondimensional bacterial and carbon concentration are *b*_0_ = 5 *×* 10^−4^ and *c*_*a*_(*t* = 0) = 10, other parameters are from table 1.

### Remotely cross-feeding populations

Next, we consider auxotrophic populations in separate reservoirs, exchanging limiting metabolites through a connecting channel. As mentioned earlier, we apply the model to an algal-bacterial system, obtaining our parameters from experiments where the phototrophic alga *L. rostrata*, auxotrophic for vitamin B_12_, is grown in co-culture with the heterotrophic bacterium *M. loti*. The algal and bacterial populations in their reservoirs have initial concentrations, *a*_0_ and *b*_0_, respectively. Neither carbon source nor vitamin (the limiting metabolites) are initially present in the reservoirs and channel. The coexistence diagrams in Figure 4a,b show what values in the initial concentration parameter space give rise to long-term mutualistic coexistence or a population crash due to metabolite deprivation. These fates are the possible fixed points of our model, which we shall also refer to as model equilibria. Figure 4a displays the boundary between these two regions for different values of the channel length *λ* for a fixed value of the equilibration length *η*. In Figure 4b crash-coexistence boundaries are instead shown for different equilibration lengths *η* at fixed *λ*. Also shown on both diagrams is the *membrane limit* (bottom-left grey line). In this limit the distance between reservoirs vanishes (*λ* → 0) and they are simply separated by a membrane impervious to cells, as has been demonstrated experimentally in co-culturing/metabolomic experiments (Paul et al. 2013). We assume instantaneous equilibration of metabolite concentrations across the membrane in this limit. It is thus an ideal case in which exchanges are not limited by diffusion dynamics along the tube nor by the geometry of the problem, and as such represents an interesting common reference case to understand the impact of both the channel length *λ* and the equilibration length *η*.

**Figure 4:**
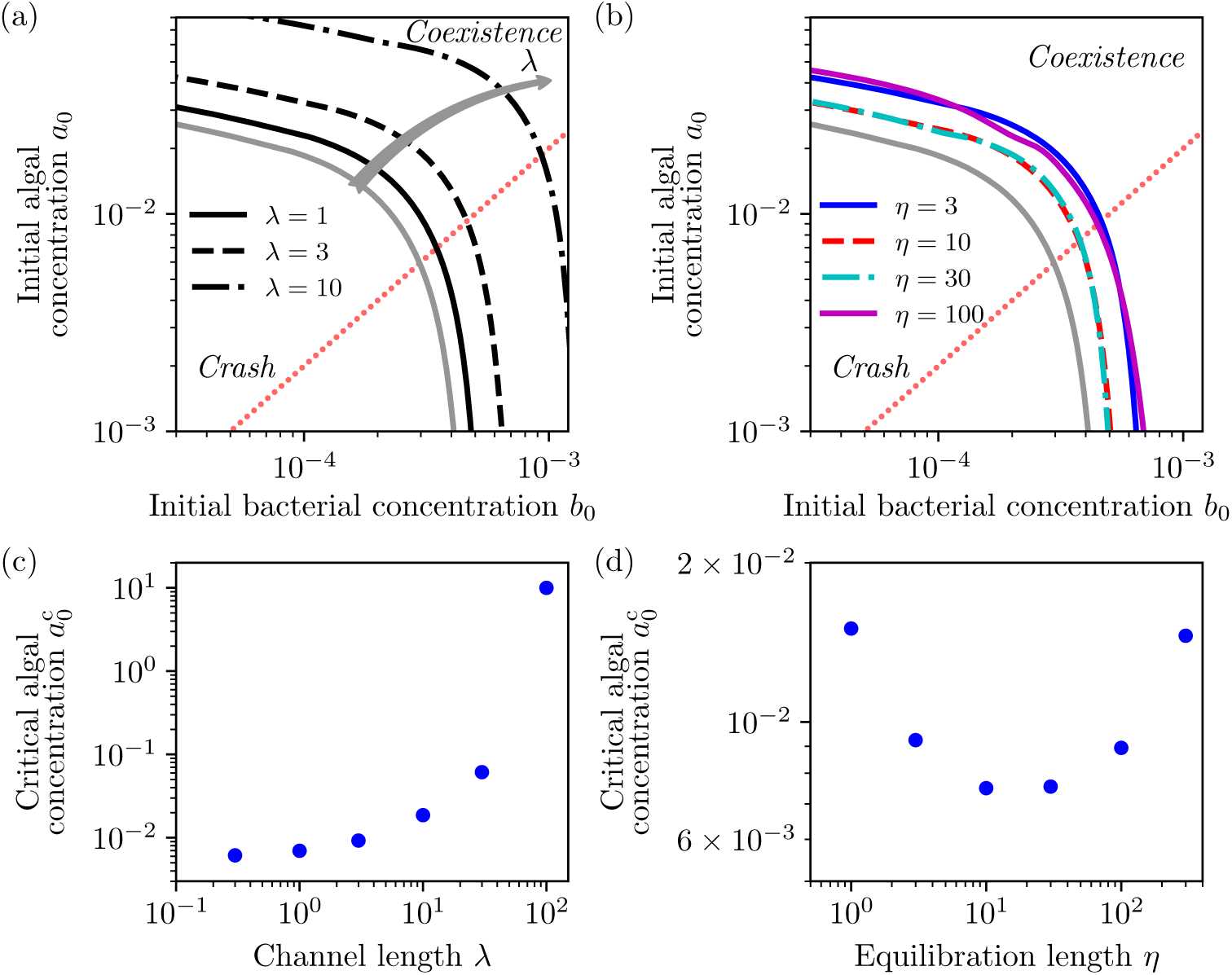
Coexistence diagram illustrating the long-time fate of mutualistic populations in terms of initial concentrations. (a) At a fixed equilibration length *η* = 3, increasing channel length *λ* causes the coexistence region to shrink progressively. (b) On the other hand, the response to an increase in *η* for fixed channel length *λ* = 3 is nonmonotonic. The coexistence initially contracts, then expands, and finally contracts again. The grey lines in both plots corresponds to the membrane limit for which *λ* → 0 and equilibration of metabolite concentrations between the two flasks is instantaneous. This provides the maximum possible concentration parameter space for mutualistic coexistence. The coexistence boundaries were determined by solving equations (2.14) and (2.15) numerically using the parameters in table 1. (c) Along the transect (dotted red line) in (a) corresponding to a conserved ratio of initial concentrations *b*_0_*/a*_0_ = 20.0, the critical initial algal concentration 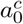 above which coexistence occurs is an increasing monotonic function of the length of tube *λ*. (d) Using the same transect in (b), the non-monotonic behavior of the critical algal concentration 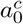 with *η* is clearly revealed.

We see that increasing the channel length has the effect of pushing the crash-coexistence boundary toward higher initial microbial concentrations (Figure 4a,c). Coexistence is achieved in the membrane limit for initial concentrations lower than those for finite *λ*. The boundary between crash and coexistence regions shifts quantitatively with *λ*, but does not change significantly qualitatively. Its shape is revealing: if the initial concentration of bacteria *b*_0_ is not too large, coexistence depends weakly on *b*_0_, and very strongly on the initial algal concentration *a*_0_. For low enough bacterial concentrations, the smallest critical initial algal concentration for which coexistence will occur increases with *λ*. These features are reasonable considering that there is a diffusive delay in the metabolite exchange between reservoirs: if the delay is too long, auxotrophs will difficultly recover in the absence of a limiting nutrient. However, we note that the model does not predict any critical length above which recovery is impossible: longer separations will simply restrict the establishment of co-existence to cases with very high initial populations.

The effect of the reservoir equilibration length *η* on the coexistence diagrams is more subtle. Recall *η* is the nondimensional ratio of growing volume to metabolite exchange area, which controls diffusive equilibration in the reservoirs. For small *η*, the crash-coexistence boundary sits above the membrane limit boundary toward higher initial concentrations. This boundary is then pushed toward lower initial concentrations for intermediate values of *η* while still sitting above the membrane limit (as expected given that the membrane limit corresponds to the ideal case of instantaneous equilibration for no separation length), before raising to higher initial values for high values of *η* (Figure 4b,d). The general shape of the boundary is preserved for all *η*. To understand the nonmonotonic dependence of the boundary shift with *η*, we note *η/λ* is the reservoir/channel volume ratio. Thus, with *λ* fixed, changing *η* takes the populations through three regimes: i) the reservoir volume is small compared to that of the channel, *η/λ* ≪ 1; ii) the volumes are the same size, *η/λ* ~ 1; iii) the channel volume is smaller than that of the reservoir, *η/λ* ≫ 1. In regime i), the equilibration time *t*_eq_ = *λη* is small, but a large channel volume relative to the reservoirs dilutes any metabolite produced, making metabolites inaccessible to the microbial partner and preventing co-existence. In regime iii), the relative channel volume is small, but co-existence is impeded due to the long equilibration time *t*_eq_ ≫ 1, which slows down significant metabolite exchanges between reservoirs. Finally, in regime ii), where reservoirs and channel have similar volume and *t*_eq_ *~* 1, mutualistic coexistence is favoured.

Aside from the co-existence or crash fixed points just discussed, we can use the model to analyse the transient dynamics leading to these equilibria. In particular, it is illuminating to evaluate the relaxation time taken for remote populations to reach the fixed points for a given initial microbial concentration in reservoirs assumed initially devoid of metabolites, as previously. Numerical solutions of the model equations show that this time varies as *λ* is increased across the co-existence/crash boundary for given *η*, as shown in Figure 5a. It is clear that the time to relax to the equilibrium rises sharply on either side of the critical *λ* at the boundary. This slow relaxation for *λ* values close to the bifurcation between extinction or co-existence is accompanied by oscillatory transients (see Figure A.4). Similar considerations apply to the dependence of this time on the equilibration length *η* for a given *λ*, within that case there is the possibility of two boundaries between extinction and survival, see figure 5b. We thus predict a complex behavior of the time needed to reach steady-state in such connected mutualistic systems, with the potential for slow relaxation if geometrical parameters are close to critical values between extinction and co-existence.

**Figure 5:**
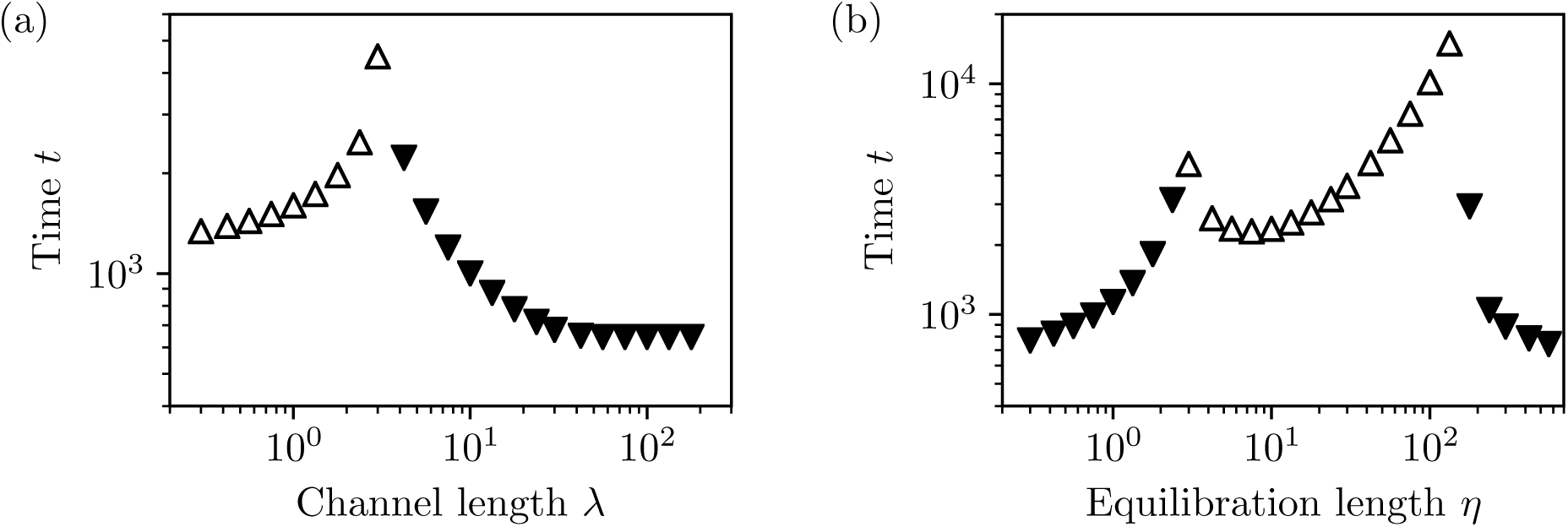
The time taken for the populations to relax to equilibrium (crash or coexistence) depends on the geometric parameters *λ* and *η*. Here, we plot these times for fixed initial microbial concentrations (*a*_0_, *b*_0_) (assuming, as before, no initial metabolites within the diffusion geometry). Times for populations reaching coexistence are shown in white up-pointing triangles, and those for populations that will crash in black down-pointing triangles. (a) For fixed *η* = 3, the relaxation time increases with *λ* up to the critical value at the coexistence boundary (where it diverges). On the other side of this critical value it decreases. (b) For fixed *λ* = 3, the dependence of the time as a function of *η* shows a similar divergence when approaching a transition between extinction and survival. For the initial concentrations (*a*_0_, *b*_0_) here chosen, two of these transitions are possible, with extinction for low and high values of *η* and coexistence for intermediate values. Both panels correspond to *a*_0_ = 2 × 10^−2^ and *b*_0_ = 3 × 10^−4^.

Interestingly, the algal and bacterial concentration fixed points, *a*, b\** respectively, are independent of *λ* and *η*, as already mentioned (see equations (A.7)). Larger separation (increasing *λ*) or weaker diffusive coupling to the reservoirs (increasing *η*) increases delays in chemical exchanges and reduces the extent of the mutualistic co-existence region. However, these geometric changes do not alter the microbial concentration fixed points, which have the same values as in the membrane limit: high densities of mutualistic microbes can be achieved even with weak or slow diffusive coupling. This equilibration is possible thanks to supply of metabolites (whose concentrations are also geometry-independent, see equations (A.7)) from the partner reservoir. A sufficiently large metabolite gradient across the channel is required to support the equilibrium metabolite and cell concentrations. Indeed, the model predicts an increase in the metabolite concentration at the production reservoir. For example, if the equilibrium concentration of vitamin B_12_ in the algal reservoir is 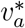, then at the bacterial reservoir we predict 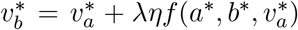, where the function *f* can be obtained by comparison with equation (A.7). The same applies for carbon. This metabolite enrichment is an interesting prediction of the model. The concentration excess at the production reservoir is linear in both separation *λ* and equilibration length *η*: two parameters with which enrichment could be experimentally controlled. As an example, for the *L. rostrata* and *M. loti* mutualism using *λ* = 1.25 and *η* = 2 (all other parameters as before) our model predicts a sevenfold enrichment of vitamin B_12_ in the bacterial reservoir compared to the algal side.

## 3 Discussion

Microbial populations often interact by diffusive exchange of metabolites in structured environments, such as the porous matrix of soil. Metabolite diffusion is known to play an important role in determining microbial dynamics in unstructured environments (Allen et al. 2013, Momeni et al. 2013, Harcombe et al. 2014, Hom and Murray 2014, Nadell et al. 2016). Current models of microbial interactions, however, do not explicitly model diffusive transport in geometrically confining habitats. A recent theoretical study has investigated microbial invasion in soil networks (Pérez-Reche et al. 2012), but interactions were modelled stochastically, without considering diffusive exchanges. How the geometry of diffusive exchanges constrains microbial interactions remains an important open question. We have addressed this here by modelling a minimal geometrical unit of microbial interaction: two mutualistc populations in finite volume reservoirs linked by a diffusive channel. The model was solved to predict the diffusively mediated interactions of mutualitistic algae and bacteria, whose dynamics in co-culture have been experimentally characterised (Kazamia et al. 2012). Two key geometrical parameters control the diffusive exchange of metabolites between the populations: the separation *λ* (the nondimensional channel length) and the equilibration length *η* (the nondimensional ratio of growing volume to metabolite exchange area). Model solutions allow prediction of whether initial concentrations of algae and bacteria will result in mutualistic coexistence or population crash (the model equilibria) for given values of the geometrical parameters *λ* and *η*. In particular, we can draw the boundary between regions exhibiting these two equilibria for given initial microbial concentration, and predict how this boundary shifts when the values of the geometrical parameters are changed.

The model makes several interesting predictions. For instance, coexistence between mutualistic partners can be achieved only if the numbers of one or both partners are abundant; low initial numbers will lead to a crash. This feature is qualitatively independent of diffusive geometry (*λ* or *η*), like the shape of the coexistence boundary itself (approximately flat for a broad range of bacterial concentrations, falling very rapidly thereafter, see figure 4). It has an intuitive explanation: an initially high concentration of one of the two species will produce a large initial amount of metabolite, which allows the partner species to grow and recover, even from initially very low numbers. A more surprising result is that mutualistic populations at a distance can achieve as high a steady concentration as in a mixed environment. The effect of the diffusive geometry is only to modify the transient dynamics and raise the initial cell concentration values required to avoid a crash. The fact that, given enough time, separated cross-feeding mutualists might reach as high numbers as populations in proximity is a counterintuitive of great potential significance for microbial ecology. This contrasts with the case of a population feeding from a distant passive resource (figure 3), for which maximum achievable concentrations do depend strongly on geometric coupling.

A final prediction of the model to highlight is the nonmonotonic dependence of the boundary position as the equilibration length *η* is varied. As one might expect, increasing the channel length *λ* (at fixed equilibration length *η* and bacterial concentration *b*_0_) increases the critical concentration of algae that will support co-existence with bacteria. On the other hand (for fixed *λ* and *b*_0_) the critical algal concentration varies nonmonotonically, falling and then rising again with increasing *η*. The dependence on *λ* is intuitive: separating the partners further increases a diffusive delay, which we recall scales like *λ*^2^, so that more algae are required to support coexistence at a distance. The nonmonotonic behaviour with *η* is less obvious. It results from a dilution of metabolites in the volume of the channel for low values of *η*, requiring higher initial densities for successful coexistence, and from weak fluxes of metabolites into the homogenisation volume when *η* is large. With respect to these two extremes, coexistence is more easily achieved at intermediate values of *η*. This is another counterintuitive prediction, which highlights the value of explicitly accounting for diffusive transport in modelling mutualistic interactions.

Our findings have implications for the microbial ecology of synthetic consortia. This is an active area of investigation, with several recent studies on microbial mutualisms (Kim et al. 2008, Kazamia et al. 2012, Allen et al. 2013, Harcombe et al. 2014, Hom and Murray 2014, Wang et al. 2014, Segev et al. 2016, McCully et al. 2017, LaSarre et al. 2017). None thus far have addressed the role of diffusive geometry on these interactions, which could test the predictions of our model. A preliminary experiment in which batch cultures of algae and bacteria grow linked by a channel allowing metabolite diffusion (filled with a hydrogel to prevent cross-contamination) demonstrates the possibility of establishing remote mutualisms, see Supporting Information. Further, it provides preliminary confirmation that vitamins accumulate in the B_12_ producer (bacteria) flask, as predicted by our model (equation (A.7)). The experiment provides a ‘proof of concept’ and a blueprint for further experiments using our connected flask set-up. These should explore how the population behaviour varies with the geometrical parameters, and if the stark predictions of the model, such as the nonmonotonicity of the crash-coexitence boundary with *η*, are borne out experimentally. Alternatively, experiments using diffusively coupled microfluidic chambers [e.g. as in the work of Kim et al. (2008)], could be used, noting that modifications would be necessary to account for stochastic effects associated with the small cell numbers in such systems (Khatri et al. 2012). As well as being tested, the model could be used to describe other synthetic consortia in which populations also interact diffusively across porous hydrogels (Kazamia et al. 2012, Harcombe et al. 2014) or microfluidic structures (Kim et al. 2008). It is straightforward to extend the model to account for two or three-dimensional diffusive exchanges appropriate to these systems.

The present model may also provide the foundation for a physical description of microbial networks, e.g. consortia for cooperative biosynthesis (Hom et al. 2015, Cavaliere et al. 2017) or microbial communities in soil, or spatially coupled biofilms (Liu et al. 2017). Indeed, as mentioned earlier, at the microbial scale, soil can be approximated as a physical network of growth chambers linked by channels (Pérez-Reche et al. 2012). In establishing the key geometric parameters that govern the most elementary unit in a network, namely two diffusively linked nodes (reservoirs), the present work provides a basis for describing population dynamics in a two- or three-dimensional network of coupled nodes (Figure 1). It is left to future work to take up the significant challenge of studying such networks, particularly when there is inhomogeneity in the diffusive couplings and stochasticity in the populations themselves. This view of microbial networks centering on the physics of diffusion could also help refine interaction matrix models of microbial communities and extend them beyond contact interactions (Mathiesen et al. 2011). An interesting possibility is that interaction networks could be simplified by constraints deriving from diffusion geometry.

Aside from the microbial networks mentioned above, the model may also be a relevant interpretative tool to understand the behaviour of structured environmental communities with diffusive exchanges, such as river biofilms (Battin et al. 2016) or sediment layers (Pagaling et al. 2014). Moreover, knowledge of the mechanisms for metabolite exchange between spatially separated organisms is important to gain insight into how such communities initiate in the natural environment, and the drivers and constraints on the evolution of mutualisms within them (Kazamia et al. 2016).

## Acknowledgments

We thank J. Kotar and R. Bowman for discussions. We thank the Cavendish and G. K. Batchelor Laboratory workshops for assistance, in particular D. Page-Croft. F.J. Peaudecerf gratefully acknowledges support from Mines ParisTech and from a Raymond and Beverly Sackler Scholarship. O.A. Croze, M.A. Bees and A.G. Smith gratefully acknowledge support from the Engineering and Physical Sciences Research Council (EP/J004847/1). O.A. Croze also acknowledges support from a Royal Society Research Grant and the Winton Programme for the Physics of Sustainability. R.E. Goldstein acknowledges support from an EPSRC Established Career Fellowship (EP/M017982/1) and the Schlumberger Chair Fund. V. Bhardwaj was in receipt of a studentship from the Gates Cambridge Trust. F. Bunbury is in receipt of a studentship from the UK Biotechnology and Biological Sciences Research Council (BBSRC) Doctoral Training Partnership.

## A SUPPORTING INFORMATION

### A.1 Diffusive reservoir equilibration (no microbes)

We consider here the purely physical equilibration between two diffusively connected reservoirs to reveal the interplay between the diffusive time and the equilibration time in such a system. This setup utilises the same geometry as in Fig. 2, with the reservoir at 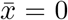 having an initial concentration 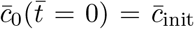 of a chemical species, and the reservoir at 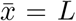 having an initial concentration 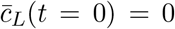 of the same species. The chemical concentration along the tube is initially equal to zero, and has diffusivity *D*. Since our focus here is purely on the different physical timescales independent of biological processes, we choose a non-dimensionalisation scheme restricted to this section only that differs from the main body of the paper. Rescaling chemical concentrations by *c*_init_, lengths by *L* and time by *L*^2^*/D*, we obtain

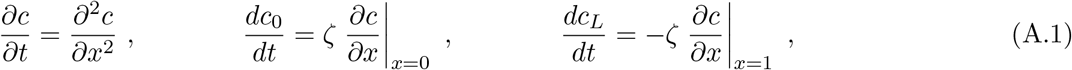

where we recognise the nondimensional parameter *ζ* = *L/ℓ*, the ratio of tube length *L* to equilibration length *ℓ* = Γ/Σ. These equations are subject to initial conditions *c*_0_(0) = 1, *c*_*L*_(0) = 0, *c*(*x*, 0) = 0 and boundary conditions *c*_0_(*t*) = *c*(0, *t*) and *c*_*L*_(*t*) = *c*(1, *t*). Despite the fact that this is a linear PDE with apparently simple boundary conditions, the fact that it exists on a finite domain, and is coupled to the reservoir dynamics, makes it difficult to obtain an explicit analytical solution for general values of *ζ*.

#### A.1.1 Approximate solution for *ζ* « 1

When *ζ* ≪ 1, the time evolution of the reservoir concentrations is much slower than the establishment of a concentration gradient in the tube. Thus, the diffusive dynamics within the tube reach a quasi-steady-state distribution between the two reservoir concentrations *c*_0_(*t*) and *c*_*L*_(*t*). In this approximation, the solution to the diffusion equation in the tube is the linear profile *c*(*x, t*) ≈ [*c*_*L*_(*t*) − *c*_0_(*t*)]*x*. Substituting this solution into the reservoir dynamics, and solving the resulting two ODEs yields (in dimensional units)

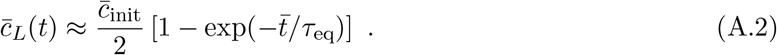

We thus deduce that in the limit *ζ* = *L/ℓ* ≪ 1, the timescale of exchanges is purely dominated by the equilibration time *τ*_eq_ = *Lℓ*/2*D*, as argued previously. The same time scale plays a role when the biological dynamics of growth and production are considered, as discussed in the main text.

#### A.1.2 General solution from Laplace transform

To find the general solution of this problem, we examine the Laplace transforms of the nondimensional concentrations 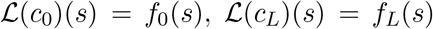, and 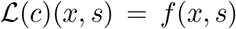. Laplace transforming the diffusion equation in the tube we find the general solution

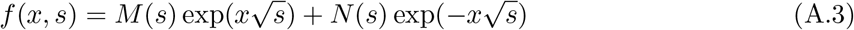

with *M*(*s*) and *N*(*s*) functions of the Laplace variable to be determined. Imposing boundary conditions at the tube ends gives

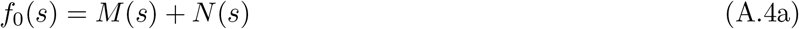

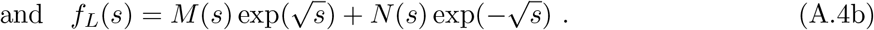

Finally, Laplace transforming the dynamical equations for the reservoir concentrations yields

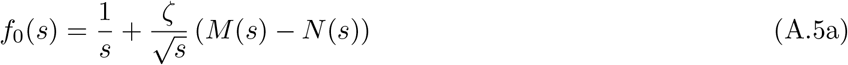

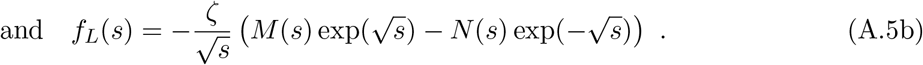

Combining the above we obtain explicit solutions for *M* (*s*) and *N* (*s*), thus entirely determining the solutions *f*_0_(*s*), *f*_*L*_(*s*) and *f*(*s*) to the problem in the Laplace space. In particular, for the concentration in the reservoir initially devoid of chemical, we obtain

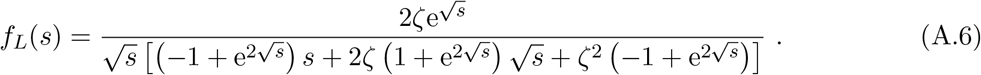

This solution in Laplace space is not easily inverted into an analytical expression for the evolution in time of 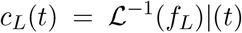. In order to access its time evolution, we adapted a numerical inverse Laplace code in Python (Barbuto 2002) which implements the Zakian method (Halsted and Brown 1972, Abate and Whitt 2006). The numerical evaluation of *c*_*L*_(*t*), as a function of the characterisic nondimensional parameter *ζ* = *L/ℓ*, is shown in figure A.1. It reveals the typical nondimensional time-scale of equilibration 1/2*ζ*, which in dimensional form becomes the previously discussed equilibration time *τ*_eq_ = *Lℓ*/2*D*. At steady state, the concentration equilibrates between the two reservoirs and the tube at a final uniform value *c*_f_ = 1/(2 + *ζ*). Finally, for *ζ* ≪ 1, the validity of the approximations of the concentration *c*_*L*_(*t*) as a saturating exponential in equation (A.2) is clearly demonstrated (Figure A.1, right panel).

**Figure A.1:**
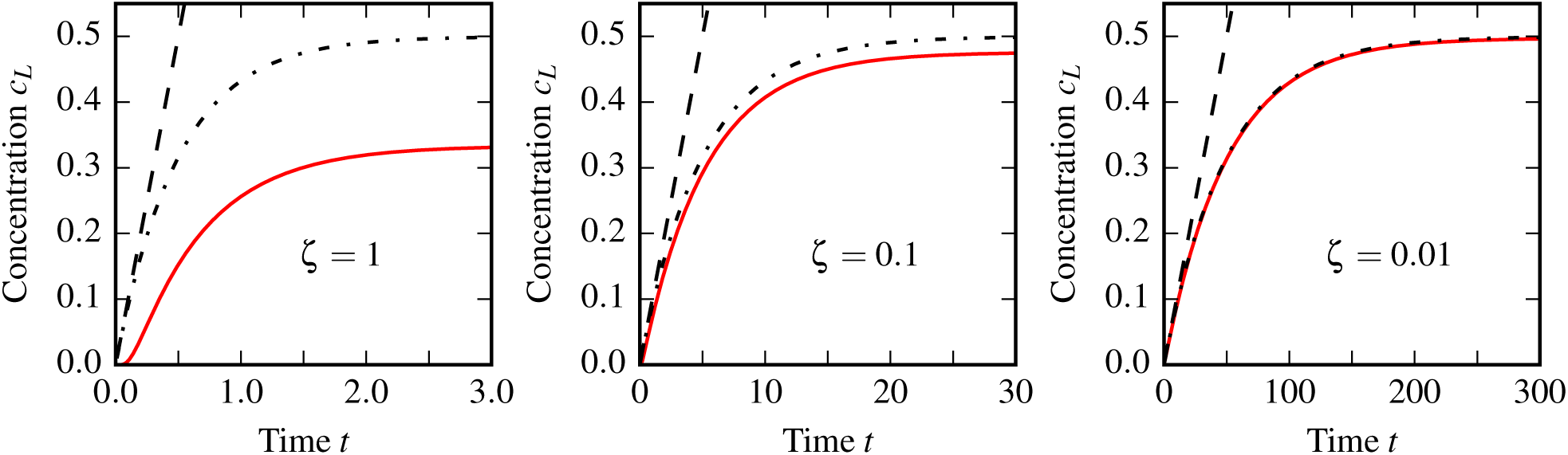
Evolution of the concentration *c*_*L*_, in a reservoir initially devoid of chemical, diffusively coupled to a reservoir filled with initial concentration *c*_0_(0) = 1. The concentration was evaluate numerically from the inverse Laplace transform of *f*_*L*_, given in equation (A.6). Each red curve corresponds to a numerical evaluation for the displayed value of the parameter *ζ*. Dash-dotted lines are the corresponding nondimensional versions of the approximation of *c*_*L*_ as a saturating exponential as given in equation (A.2), while dashed lines correspond to the linear approximation *c*_*L*_ = *ζt*. Note the change of scale of the time axis for different values of *ζ*, where time itself has been rescaled by *L*^2^*/D*.

### A.2 Mathematical model of remote mutualistic cross-feeding

#### Fixed points

The dynamical system in equations (2.7) supports a trivial set of fixed points corresponding to a system with no cells (*a* = *b* = 0) and any residual concentrations of metabolites. The non-trivial fixed point is given by

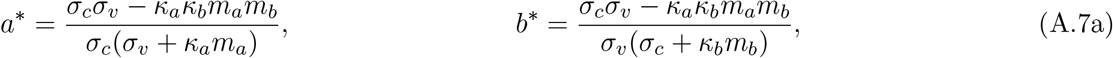

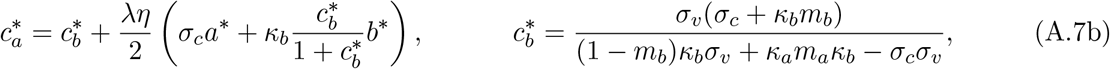

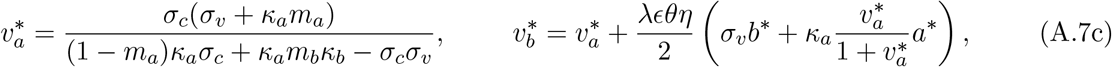

where we recall that *σ*_*j*_, with *j* = C or V, are the nondimensional metabolite production strengths; *m*_*i*_ and *κ*_*i*_ are the mortality/growth ratios and nondimensional uptake parameters for species *i* =A or B; *λ* = *L/ℓ*_*b*_ is the dimensionless channel length; *η* = *ℓ/ℓ*_*b*_ is the dimensionless reservoir volume to area ratio or equilibration length; and *ϵ* = *k*_*a*_*/k*_*b*_ and *θ* = *D*_*c*_*/D*_*v*_ are the growth rate ratio and diffusivity ratio respectively.

For the fixed point given by equations (A.7) to be physically relevant, the concentrations it describes must be positive. Therefore, the parameters must satisfy the following constraints:

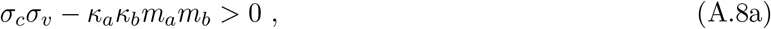

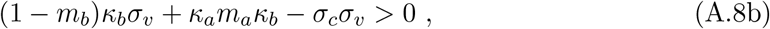

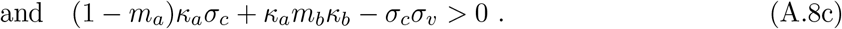

The first condition requires production strength to be strong enough to overcome cell mortality. This guarantees the existence of positive equilibrium algal and bacterial concentrations. The second and third conditions guarantee this positivity for carbon and vitamin concentrations, respectively. They require that microbial consumption be high enough to overcome production. When these conditions are satisfied, the mutualistic microbes can reach a steady-state of co-existence.

#### Membrane limit

The first natural limit of the model is that of zero channel length *λ* → 0, in which the reservoirs are in contact, but separated by a porous membrane. We call this the *membrane limit* because the membrane setup is as in membrane experiments (Paul et al. 2013), and we consider instantaneous equilibration of concentrations across the membrane as a good approximation. Fixed points for this limit are obtained trivially by letting *λ* → 0 in (A.7b)-(A.7c), which confirms that metabolite concentrations are equalised between reservoirs at steady state. We note that the membrane limit is identical to a *mixed co-culture*, where A and B grow mixed together in the same reservoir, except for the dilution effect associated with the segregation of the two species on either side of the membrane. The corresponding dynamical system for a mixed co-culture also admits a positive fixed point (*a*, b*, c*, v\**) under the same conditions (A.8), with *a\** and *b\** given by (A.7a), 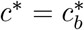 from equation (A.7b) and 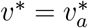 from equation (A.7c). As mentioned earlier, such a co-culture model is fundamentally different from models considering mutualistic nutrient exchanges implicitly (Murray 1989, Yukalov et al. 2012, Grant et al. 2014, Holland and DeAngelis 2010).

#### Remotely-fed monoculture

Another interesting limit is one in which a species in one of the reservoirs is replaced by a fixed concentration of metabolite. For example, we could have species B growing on C diffusing through the channel from the remote reservoir. In this limit, the model on the side of C reduces to passive diffusion from a source, which provides a useful control on the mutualistic dynamics, as mentioned in the results section. The mathematical model for such a remotely-fed monoculture is directly obtained from the remotely cross-feeding populations model (equations 2.7) by setting one microbial species and the metabolites it produces to zero.

### A.3 Parameterisation for specific microbial associations

The results presented in this paper were obtained from numerical studies of the mathematical model with parameter values corresponding to the mutualistic association between *Lobomonas rostrata*, a B_12_-requiring green alga, and *Mesorhizobium loti*, a B_12_-producing soil bacterium (Kazamia et al. 2012). The following procedure was used to obtain these parameter values. First, physiologically relevant ranges for each parameter were collected by direct measurement (see next section) or from the published literature. Then, specific parameters – both nondimensional parameters of the reduced model and dimensional parameters to convert experimental data to nondimensional units– were obtained by minimizing the squared distance between simulated time evolution, obtained through a custom finite difference solver in Python, and experimental results on mixed cultures, while searching within domains of parameter values which contain the physically relevant ones, and validating the fixed-point conditions in equation (A.8). The basin-hopping minimisation procedure gives local optima which capture well the observed dynamics of mixed co-cultures of *L. rostrata* and *M. loti* (see Figure A.2). The range of physiologically relevant parameters used to constrain the search of parameters for the association of *M. loti* and *L. rostrata* are presented in table A.1, while the fitted parameters, both dimensional and nondimensional, are given in tables A.2 and 1.

**Figure A.2:**
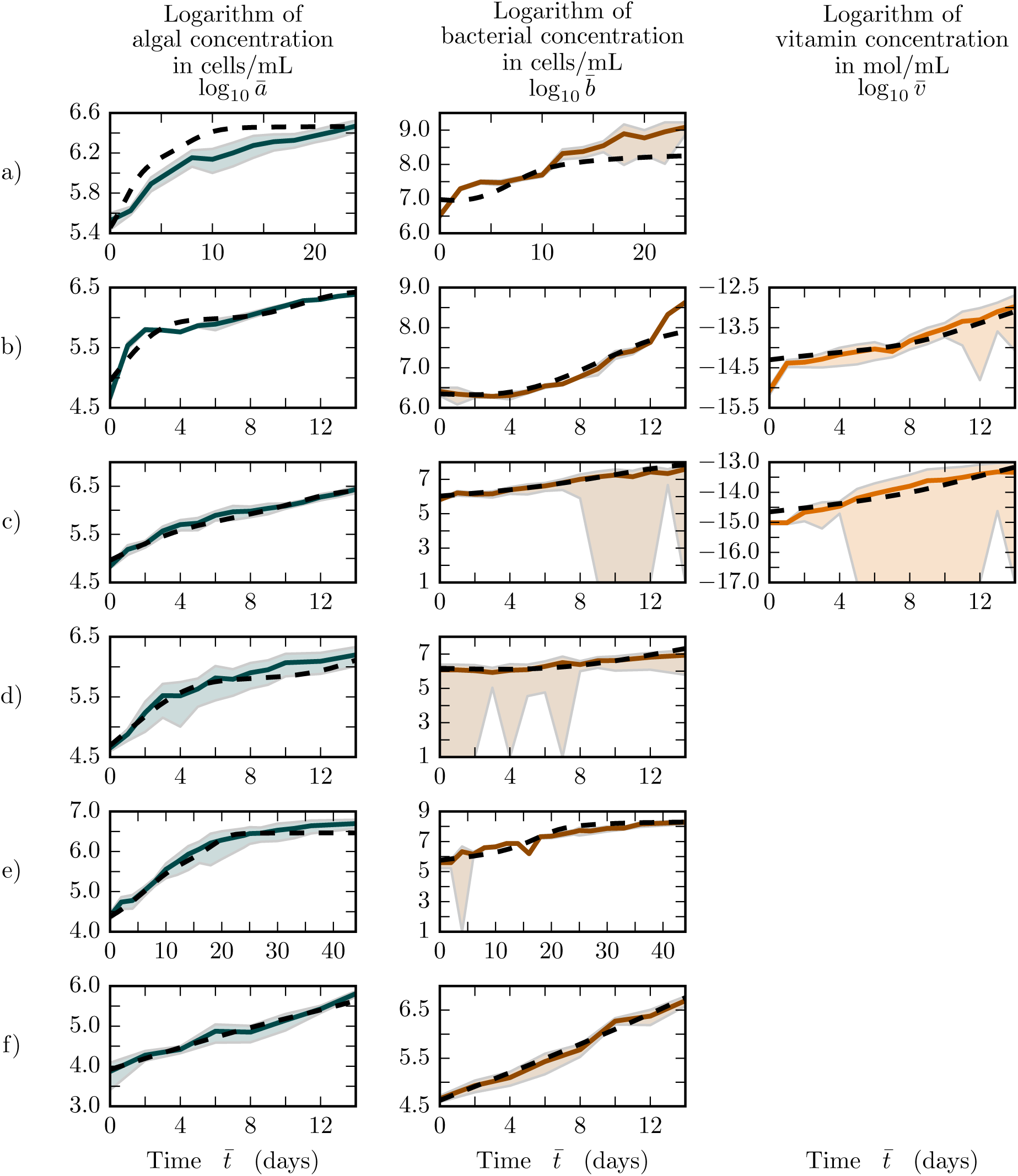
Experimental results and theoretical fits on growth of co-cultures. Rows (a)-(f) display results from 6 independent growth experiments for *M. loti* and *L. rostrata* cocultures, for different starting values and ratios of the two species. For each experiment, from left to right the panels show the algal concentration 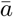, the bacterial concentration 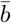, and, when data is available, the vitamin concentration 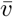. Continuous thick lines show the average value over a set of replicates, with the interval of +/- one standard deviation shown as a shaded area. The fits from the model with parameters from table 1 are shown with dashed black lines. Number of replicates per experiment from a to f is n = 6, 3, 5, 5, 4 and 4. Large downward shaded areas represent on this logarithmic scale time points for which standard deviation is comparable to the mean.

**Table A.1:**
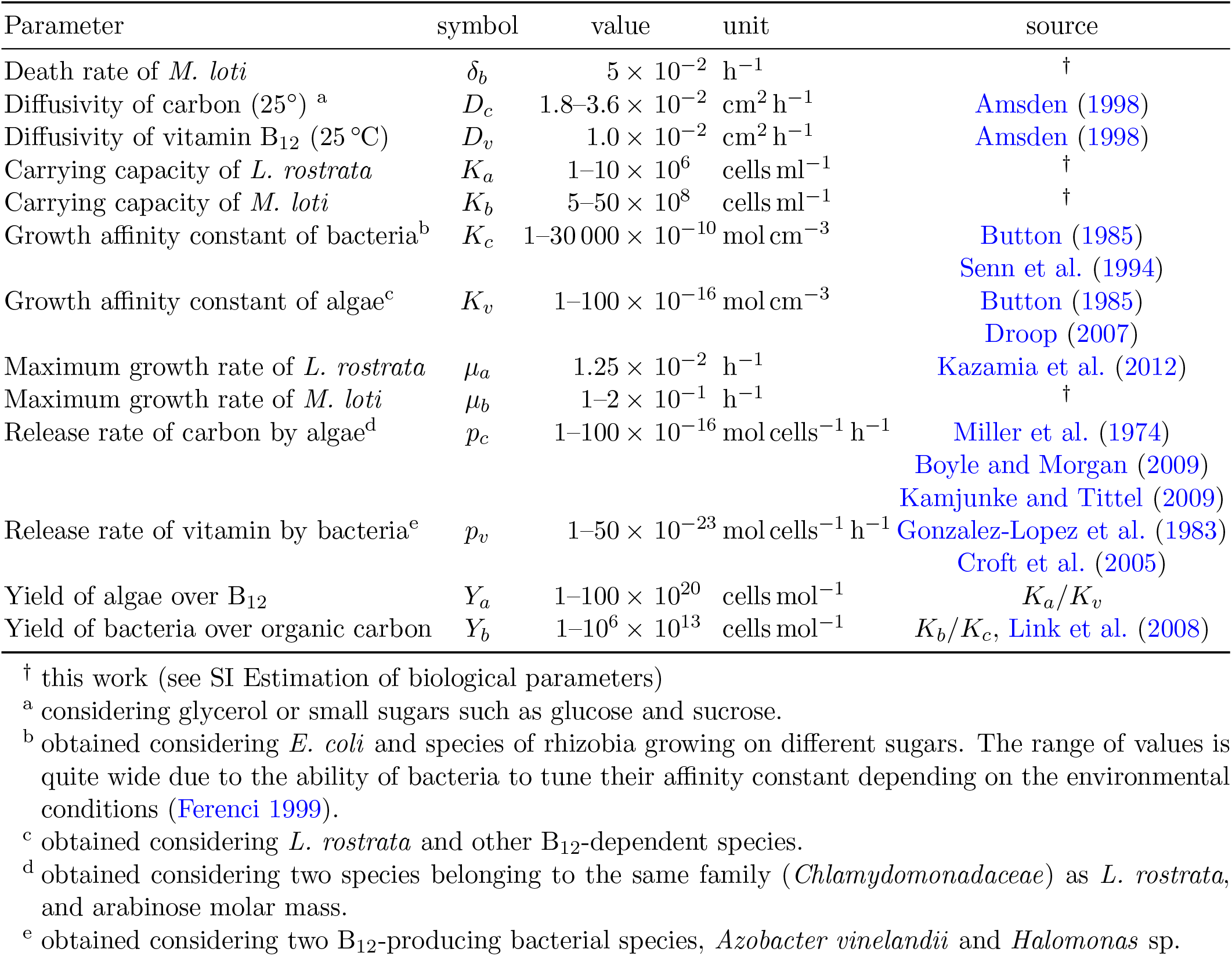
Physiologically relevant parameter ranges for the mutualistic association of *M. loti* and *L. rostrata*.

**Table A.2:**
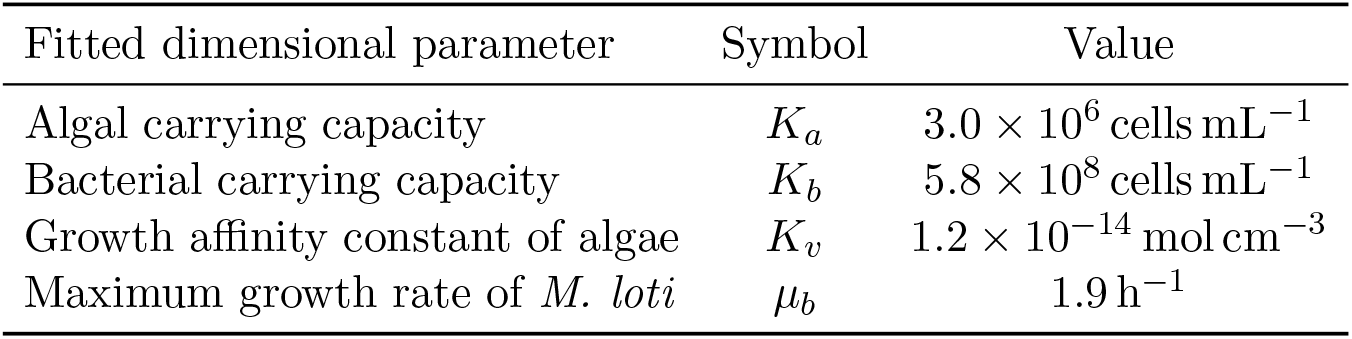
Fitted parameters for the mutualistic association of *M. loti* and *L. rostrata*.

### A.4 Estimation of biological parameters

#### Monoculture experiments: Carrying capacities of M. loti and L. rostrata

Liquid cultures of *M. loti* were grown for 3 days (33°C, shaken at 240 rpm) in TY medium (tryptone 5 g L^−1^, yeast extract 3 g L^−1^, CaCl_2_ · 2H_2_O 0.875 gL^−1^) and washed in TP+ before serial dilution for counting of colony forming units. The post-wash concentration was estimated to be 5−10 × 10^8^ cells mL^−1^. Given the existing loss of cells during washing, we therefore allow the bacterial carrying capacity of our model *K*_*b*_ to be in the range 5−50 × 10^8^ cells mL^−1^. Similarly, we estimated the carrying capacity of *L. rostrata* by growing these algae in TP+ with 100 ng L^−1^ of vitamin B_12_ for 6 days to saturation (22°C, shaken at 200 rpm, day/night cycle of 14h/10h), and plating them after washing in TP+ and serial dilution on TY agar plates for colony forming unit counting. We recorded saturation concentration *~*2 × 10^6^ cells mL^−1^, which, allowing for losses during cell washing, results in an accepted range of 1−10 *×* 10^6^ cells mL^−1^ for the algal carrying capacity *K*_*a*_ in our model.

#### Monoculture experiments: Death rate of M. loti

A pre-culture of *M. loti* in TY as above was washed in fresh TP+ and inoculated at a concentration *b*_0_ = 3.2 *×* 10^8^ cells mL^−1^ in 70 mL of TP+ without carbon source. Every two days, a 100 µL sample was taken to determine a live cell concentration through counting of colony forming units (CFUs) on TY agar. After a 2 days lag period, we measured an exponential decay of the bacterial population with death rate *δ_b_ ≈* 5 × 10^−2^ h^−1^ over the next 6 days.

#### Co-culture experiments: Global fit of model parameters

The experiments whose outcomes were used to fit the model parameters utilised the following protocol. *L. rostrata* and *M. loti* were grown in TP+ medium at 25°C on a 12h/12h day/night cycle, with 100 microeinsteins of light and shaking at 120 rpm. Bacterial concentrations were estimated with counts of CFUs on TY agar, and algal concentrations were obtained with a Coulter counter. In some experiments, B_12_ concentration was estimated with bioassays (Raux et al. 1996). Figure A.2 shows the results for a set of six independent experiments (a-f) along with global fits to the model, corresponding to the values shown in Table 1.

### A.5 Mutualism at a distance: experimental proof of concept

To test experimentally the predictions of the mathematical model, we developed a system to culture mutualistic microbial species exchanging metabolites diffusively over a finite distance. Briefly, each of two 100 mL conical Erlenmeyer flasks was modified (Soham Scientific Ltd) to have a side arm (8 mm long, outside diameter 11 mm, inside diameter 9 mm) in which a small glass tube could be inserted (25 mm long, outside diameter 8.65 mm, inside diameter 7.45 mm). Sealing of the tube-flask junction was achieved by compression of O-rings on each side of a metal washer glued onto the glass tube (see figure A.3a,b). The force of compression was established and maintained by mounting the flasks on custom sliding platforms (figure A.3b,c). To prevent contamination, flasks were capped with silicon plugs (Hirschmann Silicosen type T-22) and aluminium foil, while the middle area of the flasks and tube assembly was also further covered with aluminium foil. The central glass tube connecting the inside of both flasks was filled with a polyacrylamide (PAM) gel (4% acrylamide w/v with a relative concentration of bis-acrylamide of 2.7%, filter-sterilised before pouring, BioRad). Once polymerised, the gels in their tubes were put in a bottle of sterile water and left to soak for 6 days to allow for any of the toxic non-polymerised monomer to diffuse out of the gel. We verified the very weakly hindered diffusion of B_12_ through this gel by colorimetry, measuring a reduction of *~* 10% of diffusivity with respect to B_12_ diffusion in water, which validates the chosen gel pore size as allowing the diffusive transport of small metabolites. We also performed a test to check for cross-migration of the mutualistic species. Both flasks were filled with a rich bacterial medium for soil bacteria (TY), but only one side was inoculated with *M. loti* (see below for strain details). These bacteria reached a saturation density within a few days, but over a timescale of 2.5 months no bacteria were detected in the first flask, proving the PAM gel is not penetrable by bacteria (and by inference by the algae, which are larger).

**Figure A.3:**
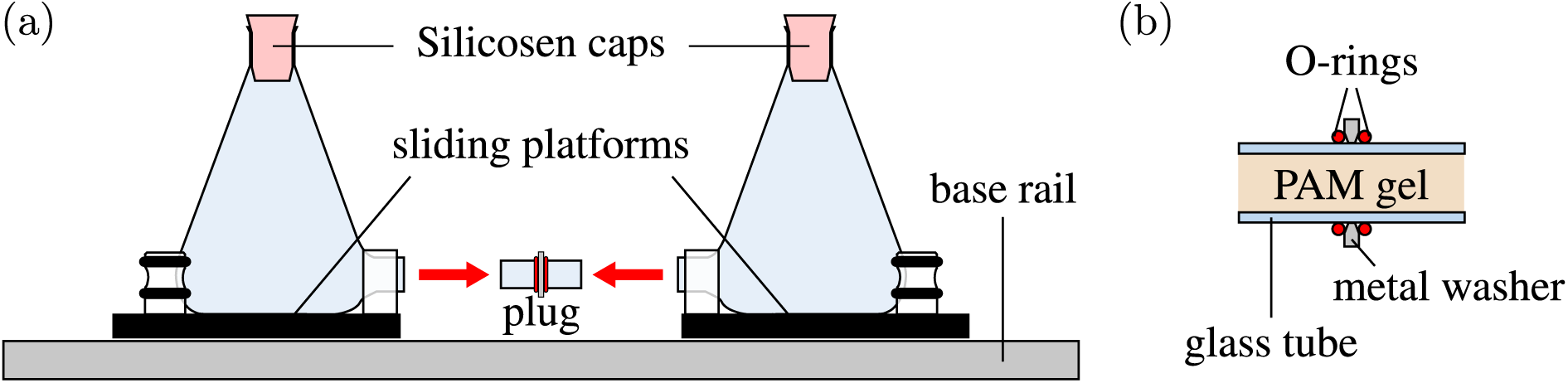
Chambers for proof-of-principle experiments. (a) Sketch of the platform holding the modified flasks during assembly. (b) Sketch of the diffusive plug filled with polyacrylamide (PAM) gel, used to connect the two flasks in experiments of mutualism at a distance.

In such connected flasks, we inoculated one side with the B_12_-dependent green alga *Lobomonas rostrata* (SAG 45-1, wild type strain) and the other with the B_12_ producing bacterium *Mesorhizobium loti* (MAFF 303099, wild type strain, original gift from Prof. Allan Downie, John Innes Centre, UK). Both inocula were diluted with TP+ medium (Kazamia et al. 2012) to the desired starting concentrations of microbes. The *L. rostrata* pre-culture was grown in TP+ with 100 ng L^−1^ of vitamin B_12_ from colonies picked from a slant, while the *M. loti* pre-culture was grown in TY medium. Both pre-cultures were washed in fresh TP+ before inoculation in the assembly in order to remove any organic carbon and B_12_ in the initial growth media. The initial concentrations of *M. loti* and *L. rostrata* were *b*_0_ = 2.2 *×* 10^8^ cells mL^−1^ and *a*_0_ = 5.3 *×* 10^4^ cells mL^−1^, inferred from viable counts. To ensure culture sterility, flask assembly and inoculation were carried out in a laminar biosafety cabinet (PURAIR VLF 48). The connected flasks were mounted on a shaking platform (120rpm) within an incubator for 50 days, at 25°C, with continuous illumination (80 µmol m^−2^ s^−1^). After this period, these assemblies were left in static incubation at 20 *±* 2°C and at ambient day/night light levels.

#### Viable counts and B_12_ concentration measurements

Algal and bacterial populations were sampled 55 and 230 days after inoculation. No contamination (external or between species) was detected, and PCR screening was used to confirm species identity as *Mesorhizobium loti* bacteria and *Lobomonas rostrata* algae. This confirms the ability of the PAM gel to prevent cells from crossing, while allowing metabolites to be exchanged.

Viable counts revealed that the population of bacteria 55 days after inoculation was *~* 10^3^ smaller than the inoculum. At the same time point the algae had grown little: the cell concentration was only 1.3 times larger than the inoculum. After 230 days the bacteria had recovered, and the algae had grown significantly. At this time the algal concentration from two replicates was *a* = 7.8 *±* 0.3 *×* 10^5^ cells/cm^3^ (where the uncertainty is the standard error in the mean), about 15 times the inoculation concentration and close to the carrying capacity they reach in well-mixed co-cultures (see table A.1). While slight initial growth of the algae might be attributed to internal reserves of vitamin B_12_, it is difficult to account for growth 230 days after inoculation in the absence of the vitamin. Indeed, using bioassays (Raux et al. 1996) we measured a B_12_ concentration of 24 *±* 3 pg/ml in the medium on the side of the algae. On the side of the bacteria, we found 132 *±* 7 pg/ml. This implies the existence of a concentration gradient across the tube between the two flasks. This is required for the supply of the B_12_ to the algae, as predicted by the model (see equation A.7c).

**Figure A.4:**
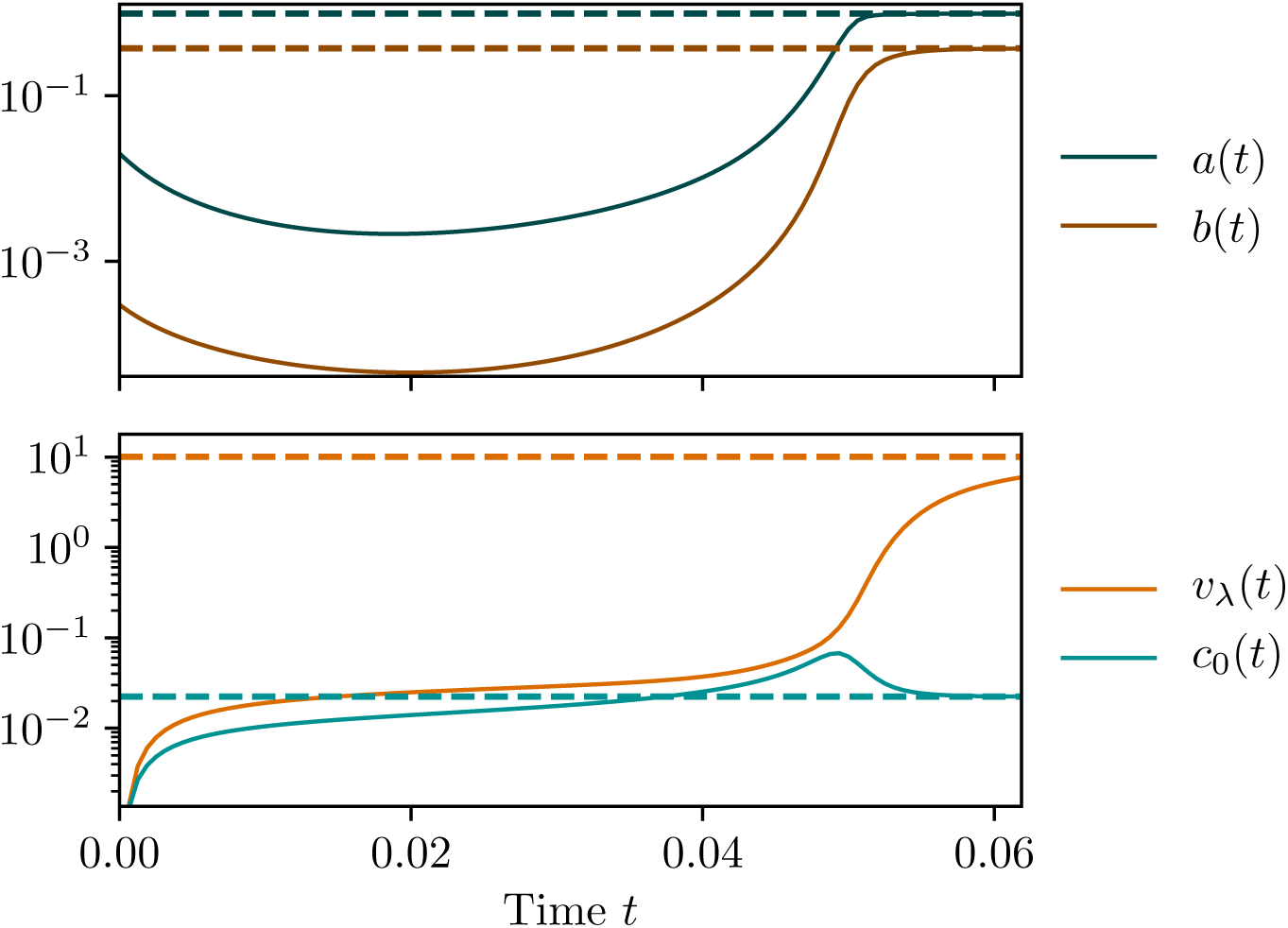
Example of oscillations of concentrations of cells and metabolites before convergence. Initial parameters are close to the boundary between survival and extinction (*λ* = 2, *η* = 3, *a*_0_ = 2 *×* 10^−2^, *b*_0_ = 3 *×* 10^−4^ and no initial nutrients).

